# Near infrared conjugated polymer nanoparticles (CPN™) for tracking cells using fluorescence and optoacoustic imaging

**DOI:** 10.1101/2023.01.30.526261

**Authors:** Ana Muñiz-García, Alejandra Hernandez Pichardo, James Littlewood, Jack Sharkey, Bettina Wilm, Hannah Peace, Dermott O’Callaghan, Mark Green, Arthur Taylor, Patricia Murray

**Affiliations:** Department of Molecular Physiology and Cell Signalling, Institute of Systems, Molecular and Integrative Biology, University of Liverpool, Liverpool, UK; Centre for Genomics and Child Health, Blizard Institute, Faculty of Medicine and Dentistry, Queen Mary University of London, London, UK; Centre for Pre-clinical Imaging, University of Liverpool, Liverpool, UK; iThera Medical GmbH, Munich, Germany; Perkin Elmer, UK; Stream Bio, Alderley Park, UK

## Abstract

Tracking the biodistribution of cell therapies is crucial for understanding their safety and efficacy. Optical imaging techniques are particularly useful for tracking cells due to their clinical translatability and potential for intra-operative use to validate cell delivery. However, there is a lack of appropriate optical probes for cell tracking. The only FDA-approved material for clinical use is indocyanine green (ICG). ICG can be used for both fluorescence and photoacoustic imaging, but is prone to photodegradation, and at higher concentrations, undergoes quenching and can adversely affect cell health. We have developed novel near-infrared imaging probes comprising conjugated polymer nanoparticles (CPNs™) that can be fine-tuned to absorb and emit light at specific wavelengths. To compare the performance of the CPNs™ with ICG for *in vivo* cell tracking, labelled mesenchymal stromal cells (MSCs) were injected subcutaneously in mice and detected using fluorescence imaging (FI) and a form of photoacoustic imaging called multispectral optoacoustic tomography (MSOT). MSCs labelled with either ICG or CPN™ 770 could be detected with FI, but only CPN™ 770-labelled MSCs could be detected with MSOT. These results show that CPNs™ show great promise for tracking cells *in vivo* using optical imaging techniques, and for some applications, out-perform ICG.

## Introduction

Cell therapies have potential for treating various conditions, including cancer,^1^ degenerative diseases ^2^ and acute tissue injury.^3^ One of the barriers facing the development and optimisation of these therapies is that it can be difficult to track the cells *in vivo* following their administration. Without knowing the biodistribution and fate of the cells, their safety and efficacy cannot be adequately assessed. Tracking cells *in vivo* can also shed light on their mechanisms of action. For instance, using bioluminescence imaging (BLI) to assess the biodistribution of kidney-derived regenerative cells and mesenchymal stromal cells (MSCs), we previously found that following intravenous (IV) administration in mice, the cells were entrapped in the lungs and did not persist beyond 24 hours.^4-6^ This showed that the therapeutic effects of the cells were not due to them homing to target organs such as the kidney and replacing injured host cells as previously thought,^7,8^ but were instead due to paracrine or endocrine effects.^9^

Imaging cells *in vivo* with BLI requires that the administered cells express a luciferase enzyme, the most common being firefly luciferase (Fluc). In the presence of oxygen, ATP, Mg^2+^ and the substrate luciferin, Fluc catalyses the production of light. BLI is a very effective technique for tracking cell fate because in addition to indicating the location of the cells, it also shows if they are alive or not; this is because light is only emitted from Fluc-expressing (Fluc^+^) cells if they are viable.^4^ The generation of Fluc^+^ cells typically involves introducing the Fluc transgene by lentiviral transduction, then selecting the Fluc^+^ cells and expanding them in culture to generate sufficient numbers for *in vivo* administration. This can be problematic for some primary cell types, such as MSCs, which can only be passaged a certain number of times before they become senescent. Other problems with BLI include poor spatial resolution (2-5mm) and low penetration depth (∼1cm).^10^ Moreover, the requirement for a substrate to be administered means BLI is not suitable for clinical imaging.

Some of these problems can be overcome by using cell tracking nanoprobes that can rapidly label the majority of cells within a population and enable them to be imaged using a clinically translatable imaging modality such as fluorescence imaging (FI) or multispectral optoacoustic tomography (MSOT).^11^ MSOT is an emerging technology that is well-established for small animal imaging, and its effectiveness as a diagnostic tool in human patients is being assessed in a variety of clinical trials.^12^ MSOT is a type of photoacoustic imaging that involves illuminating a subject with near infrared (NIR) laser light, whereby photoabsorbers present within the tissue absorb the light and undergo thermoelastic expansion, generating acoustic waves that can be detected at the body surface. Whereas BLI and FI have poor spatial resolution (2-5 mm) and relatively low penetration depth (1-2cm), MSOT has the advantage of higher spatial resolution (∼150 µm) and greater penetration depth (4-5 cm).^10^ Moreover, in contrast to BLI and FI which are typically used to generate planar images, MSOT produces a tomographic image, allowing the position of cells to be identified in 3D.

Near infrared (NIR) nanoprobes that absorb and emit light in the “optical window” (650 nm-900 nm) are particularly suited for tracking cells with both FI and MSOT *in vivo* because within these wavelengths, there is limited absorbance by haemoglobin. Indocyanine green (ICG) is an NIR dye that has been used successfully as a contrast agent in both FI and MSOT applications in small animals,^13-15^ and is FDA-approved for various clinical applications, including the assessment of liver function and the identification of tumour margins during surgery.^16,17^ ICG has also been used to label cells and track them *in vivo* in mice using FI and MSOT.^14^ However, one of the disadvantages of ICG is that it undergoes a degree of photodegradation,^18^ which affects signal intensity, and also has a tendency to leach out of the labelled cells,^14^ potentially labelling surrounding cells and tissues and leading to false positive results.

To overcome the problems with ICG, we developed a novel type of NIR nanoparticle called “conjugated polymer nanoparticles” (CPNs™)^19^ and assessed their potential for cell tracking. CPNs™ are next generation organic imaging agents, which exhibit exceptional emission brightness and stability whilst avoiding the use of heavy metals sometimes found in other nanoparticle systems.^20^ The particles used in the study are approximately 60 – 70 nm in diameter, with a carboxylate rich surface and contain magnetic iron oxide particles. The embedded iron oxide particles offer a further imaging modality if required (i.e., magnetic resonance imaging) and further provide the benefit of magnetic materials, such as ease of manipulation and purification via magnetic separation. Quantum yields of the materials are estimated to be ca. 1-2%,^20^ yet brightness should be significantly higher relative to other nanomaterials with similar quantum yields due to their large extinction coefficient.^21^

Here, we have assessed the potential of a range of NIR CPN™ probes (CPN™ 770, CPN™ 820, CPN™ 830, CPN™ 840, CPN™ 1000) for labelling human umbilical cord tissue-derived MSCs (hUC-MSCs). We imaged them *in vitro* with NIR fluorescence microscopy, and have compared the performance of CPN™ 770 nanoprobes with ICG for cell tracking in mice using FI and MSOT. We found that all NIR CPNs™ could be used to detect labelled hUC-MSCs with confocal microscopy without any noticeable effect on cell viability. *In vitro* analysis showed that CPN™ 770 probes displayed the highest radiant efficiency as well as the most intense signal with MSOT, indicating that these probes would likely be the most effective for tracking cells *in vivo*. Using flow cytometry, we compared the labelling efficiency of different concentrations of CPN™ 770 nanoprobes in comparison with ICG. With both ICG and CPN™ 770 (irrespective of the concentration used), the majority of hUC-MSCs within the population were labelled. To confirm that the cells remained viable *in vivo*, Fluc^+^ hUC-MSCs were labelled with either CPN™ 770 nanoprobes or ICG, and following subcutaneous injection into mice, were imaged with BLI. FI showed that the performance of CPN™ 770 and ICG was similar, with both tracking agents allowing the cells to be easily detected. However, with MSOT, ICG-labelled hUC-MSCs were barely detectable, whereas those labelled with CPN™ 770 were easily visible. The CPN™ 770 nanoprobe therefore has some advantages over ICG and shows great promise for future cell tracking applications with both FI and MSOT.

## Experimental

### Materials

Unless otherwise indicated, general reagents and cell culture reagents were purchased from Sigma-Aldrich. Conjugated polymer nanoparticles with a carboxylate functionalised surface were supplied by Stream Bio (CPN™ 770, CPN™ 820, CPN™ 830, CPN™ 840, CPN™ 1000) and used as received without further purification (batch numbers CPN770 – 20062977021115; CPN820 – 20090382021115; CPN830 – 21070883021019; CPN840 – 20081384021115; CPN1000 – 200714100021115).^19^

### Cell culture

hUC-MSCs were obtained from NHS Blood and Transplant (NHSBT, UK) after passage 2. hUC-MSCs expressing the luc2 firefly luciferase (FLuc) reporter (FLuc^+^ hUCMSCs) were used for *in vivo* experiments as a tracking control. To generate FLuc-expressing cells, lentiviral transduction was undertaken using a vector encoding the FLuc reporter and ZsGreen under the control of the constitutive elongation factor 1-α (EF1α). The pHIV-Luc-ZsGreen vector was kindly gifted by Bryan Welm and Zena Werb (Addgene plasmid #39196). ^22^ Both unmodified hUC-MSCs and FLuc^+^ hUC-MSCs were expanded in minimum essential medium α (MEMα) containing GlutaMAX (32561-029, Gibco) supplemented with 10% fetal bovine serum (FBS; 10270-106, Gibco) in the presence of 1% penicillin– streptomycin, and maintained in a regular humidified air incubator (approx. 90–95% humidity) set at 5% CO2 and 37°C.

hUC-MSCs or FLuc^+^ hUC-MSCs were labelled with a range of near infra-red (NIR) CPN™ probes (CPN™ 770, CPN™ 820, CPN™ 830, CPN™ 840, or CPN™ 1000; Stream Bio, UK). All probes were used from a stock concentration of 1×10^9^ probes/ml, diluted in fresh culture media (MEMα with 10% FBS, 1% penicillin-streptomycin), and incubated with sub-confluent cells for 24h prior to any *in vitro* analysis and/or *in vivo* administration. In parallel, cells were also labelled with 100 µg/ml ICG (27462, Cayman Chemical, 10mg/ml in DMSO) for comparison. ICG cell staining consisted of a 30 min incubation in culture media, followed by two consecutive washes with fresh medium to remove any traces of ICG in suspension. Following staining, cells were kept in regular culture medium for approximately 24h prior to any *in vitro* analysis and/or *in vivo* administration.

Cells labelled with NIR-CPN™ nanoprobes or ICG were either fixed for microscopy imaging (see below) or suspended in phosphate buffered saline (PBS) for flow cytometry (see below) and/or animal experiments. To obtain a suspension of hUC-MSCs, cells were incubated with 0.25% Trypsin-EDTA for a maximum of 5 min. Cells were counted using a TC20 Automated Cell Counter (BioRad), centrifuged at 400xg for 3 min, and suspended to the density required for each specific downstream analysis.

### Microscopy

Following labelling with CPN™ nanoprobes or ICG, cells cultured in chamber slides were fixed with 4% paraformaldehyde (PFA) in PBS for 10 min and mounted using Fluoroshield™ with DAPI (F6057, Sigma). To capture images, an Andor Dragonfly spinning disk microscope system coupled to an EMCCD camera was used with a 40x/1.3 oil objective. Images were captured using the 637 or 750 nm laser lines. The emission filters used were 600/50 or 700/75. Image visualization was done using the IMARIS version 9.9.0 (Bitplane, Schlieren, Switzerland) software package.

### Flow cytometry

Cells in suspension were transferred to flow cytometry tubes with strained caps (35 µm mesh; Corning™, FisherScientific) and 10,000 events were analysed per sample. Data were acquired on a BD CANTO II flow cytometer using BD FACSDiva software (BD Biosciences) using a 633 nm excitation laser and a 780/60 emission filter. Data analyses were performed using the FCSalyzer 0.9.22 software.

### Animal experiments

Three male eight to ten-week-old C57BL/6 albino mice were used for all animal experiments. Mice were housed in individually ventilated cages (IVCs) under a 12 h light/dark cycle and provided with standard food and water *ad libitum*. All animal procedures were performed under a license granted by the Home Office under the Animals (Scientific Procedures) Act 1986 ^23^ and were approved by the University of Liverpool Animal Welfare and Ethics Review Board. Prior to cell administration, fur was removed with clippers and depilatory cream (Veet Hair Removal Cream 8336076, RB Healthcare, UK). Mice received a total of 4 subcutaneous (SC) injections: 5×10^5^ FLuc^+^ hUC-MSCs labelled with CPN™ probes into the bottom right flank; 5×10^5^ unlabelled FLuc^+^ hUCMSCs into the bottom left flank; 100 µl of CPN™ probes stock solution into the top left flank; and 5×10^5^ FLuc^+^ hUC-MSCs labelled with ICG in the top right flank (Fig. 3A). Animals were imaged shortly after injection and the whole experiment was performed under terminal anaesthesia with isoflurane.

**Fig. 1.**
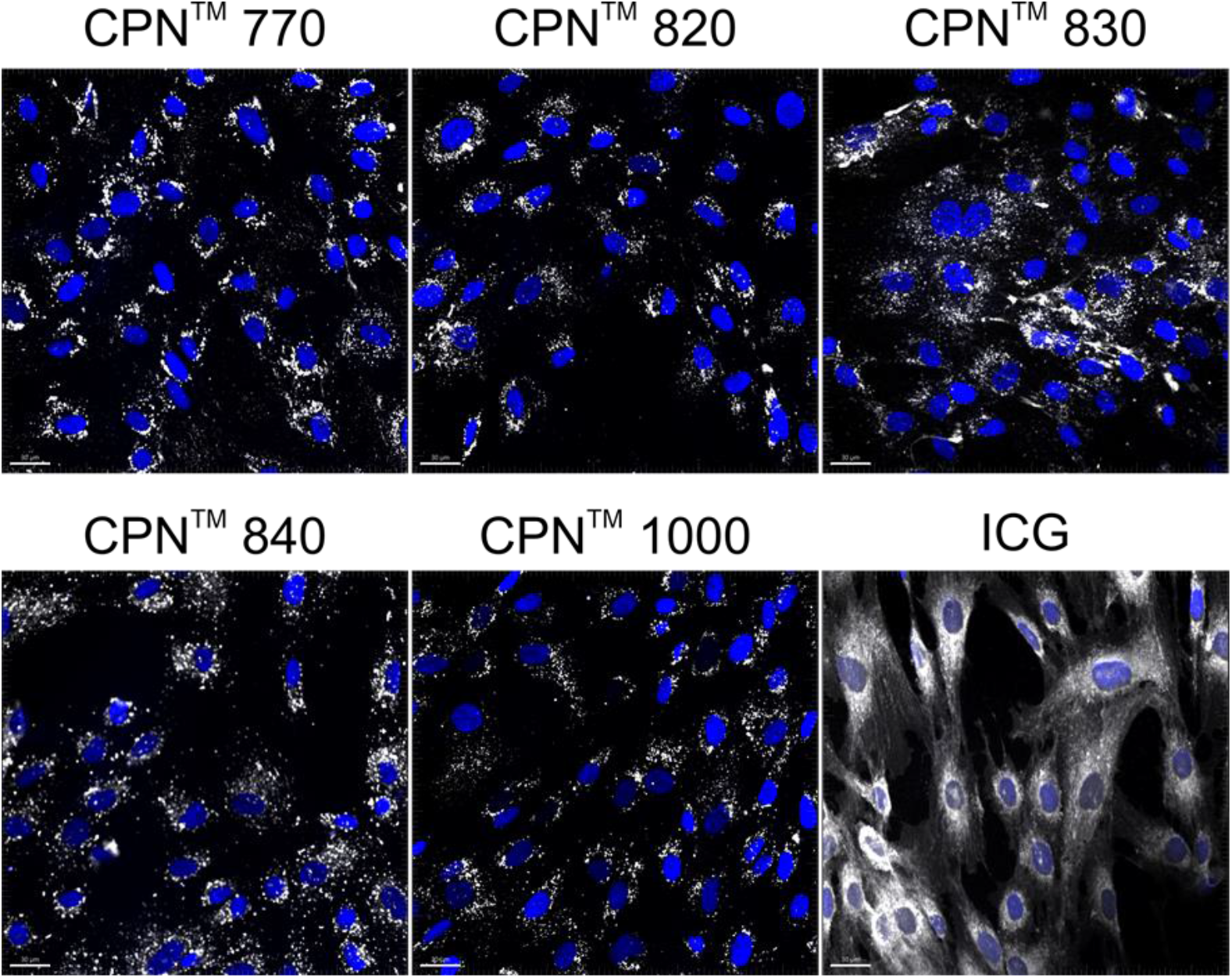
*In vitro* labelling of hUC-MSCs with NIR CPN™ nanoprobes (1×10^8^ particles/ml) and ICG (100 µg/ml). Cells were fixed 24h following labelling and imaged by confocal microscopy. Scale bar, 30µm.

**Fig. 2.**
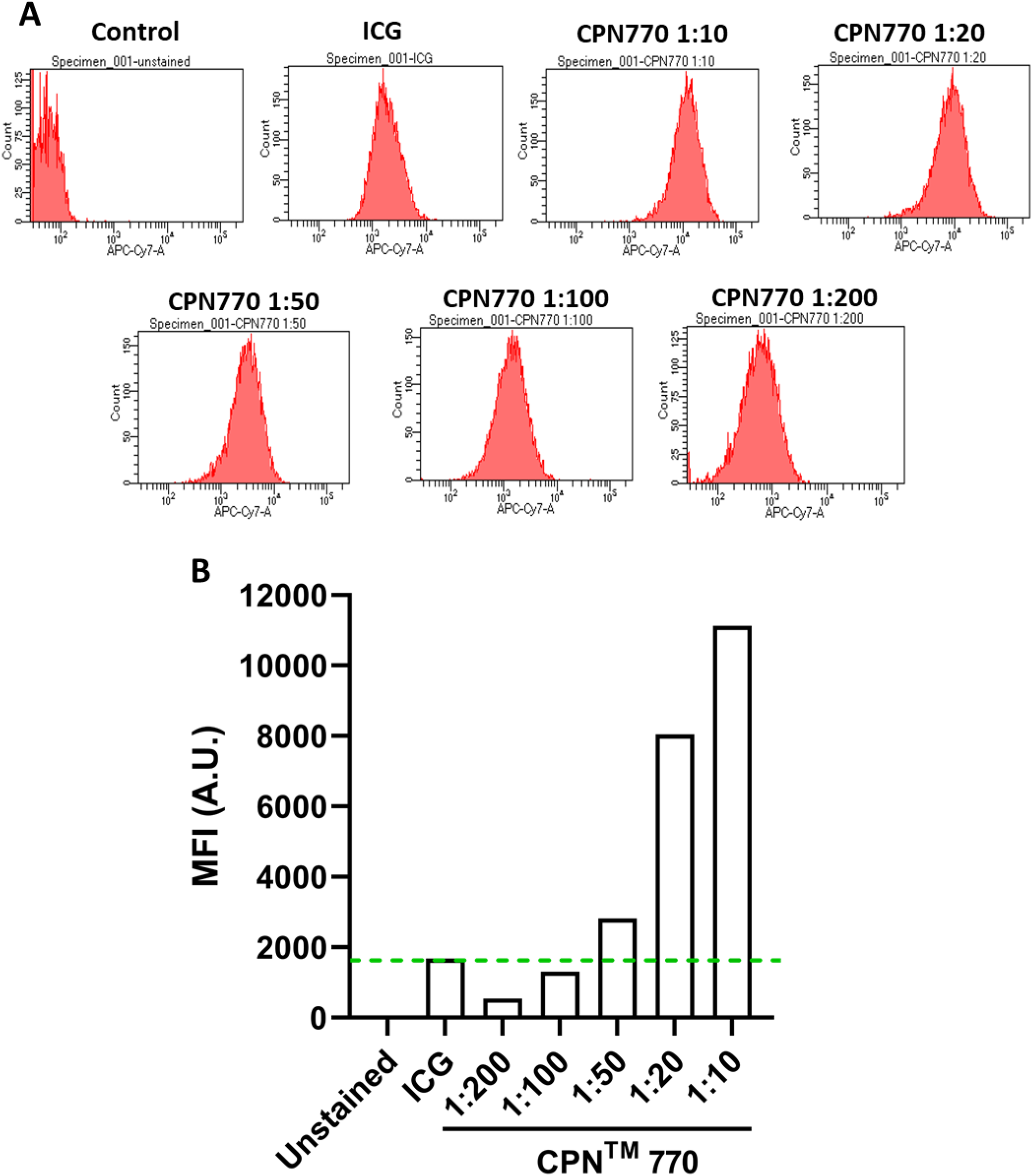
Flow cytometry of hUC-MSCs labelled with a concentration range of CPN™ 770 nanoprobes. (A) Histograms of unlabelled cells (negative control), cells labelled with 100 µg/ml ICG (positive control) or a 1:10, 1:20, 1:50, 1:100, 1:200 dilution of a stock concentration of 1×10^9^ CPN™ 770 nanoprobes/ml. (B) Mean fluorescence intensity (MFI) of the different probes in arbitrary units (A.U.).

**Fig. 3.**
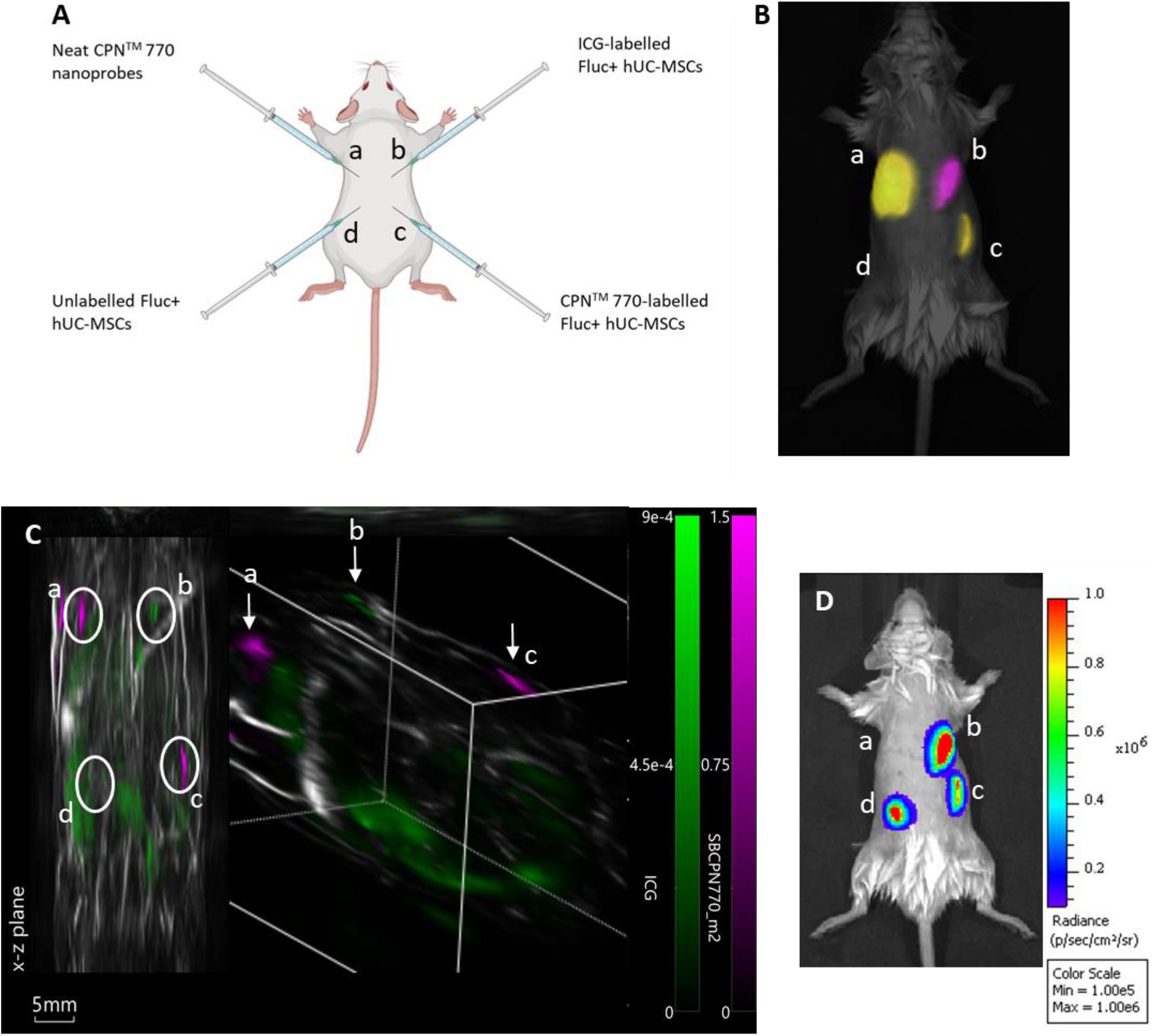
Multimodal *in vivo* imaging of Fluc^+^ hUC-MSCs labelled with CPN™ 770 nanoprobes or ICG following subcutaneous injection. (A) Schematic showing the sites of subcutaneous injection of the Fluc^+^ hUC-MSCs or neat CPN™ 770 nanoprobes. Fluc^+^ hUC-MSCs were labelled with 1×10^8^ CPN™ 770 nanoprobes/ml for 24h, or with 100 µg/ml ICG for 30mins. 0.5×10^6^ cells were injected in 100µl of saline, and neat CPN™ 770 nanoprobes were injected at a concentration of 1×10^8^ particles in 100µl. (B) Representative fluorescence image of injected mouse shortly after cell administration, after spectral unmixing. Unlabelled Fluc^+^ hUC-MSCs served as a negative control. (C) MSOT image of the same mouse; left image shows x-z plane and right image shows tomographic reconstruction, with two different colour scales for each probe. (D) Bioluminescence image of the same mouse shown in B and C. n=3.

### Fluorescence imaging (FI)

*In vivo* FI studies were performed before BLI. The mice were imaged using an IVIS imaging system (IVIS® Spectrum, Perkin Elmer) and detection was performed using a range of excitation (ex)/emission (em) filter combinations for spectral unmixing. These were 605(ex):760,780,800,820,840(em); 710(ex):780,800,820,840(em); 745(ex):820,840(em) nm. Acquisition was performed with a 13.3 cm FOV, f-stop of 2 and binning of 8. Spectral unmixing for ICG and CPN™770 was undertaken with Living Image v. 4.5.2 (Perkin Elmer) and images are shown as colour-coded composites of the two probes.

### Bioluminescence imaging (BLI)

For *in vivo* BLI, mice received a subcutaneous injection of D-Luciferin (Promega, UK) (10 μL/g [body weight] of a 47 mM stock solution) after FI. 20 min after administration of the substrate, the animals were imaged with the IVIS system. Data are displayed as radiance (photons/second/centimeter^2^/steradian), where the signal intensity scale is normalised to the acquisition conditions. Acquisition was performed without an emission filter, a 13.3 cm FOV, f-stop of 1, and a binning of 8.

### Multispectral optoacoustic tomography (MSOT)

MSOT was performed using the inVision 256-TF system (iThera Medical, Germany). Images were reconstructed in viewMSOT 4.0.1.34 software (iThera Medical, Germany) using the BP 4.0 pre-set. Reconstruction FOV was set to 25 mm. The isoflurane dose was titrated to produce a respiratory rate of 1 Hz. The mouse was placed in the MSOT mouse holder supine with a thin layer of clear non-absorbing ultrasound gel applied (Barclay-Swann, UK). The holder and mouse were transferred into the inVision 256-TF water bath previously heated to 34°C. The mouse was allowed to equilibrate to the water bath temperature for 15 minutes before imaging was started. Mice were imaged from neck to hindquarters in 1 mm steps. 10 frames were acquired per stage position and wavelength then averaged. Mice were imaged at 36 wavelengths: from 700 to 875 nm in steps of 5 nm. After image reconstruction, images were spectrally unmixed for haemoglobin, oxyhaemoglobin, ICG, and CPN™ 770 using the linear regression algorithm (viewMSOT, iThera Medical, Germany) and their a priori spectra.

## Results and discussion

### Cellular uptake of NIR CPNs™

To investigate if the NIR CPNs™ were taken up by hUC-MSCs, the cells were incubated with each type of CPN™ at a concentration of 1×10^8^ particles/ml for 24h, or with ICG at a concentration of 100 µg/ml for 30mins as previously described.^13^ This concentration of ICG was used because a previous report has shown that higher concentrations lead to a statistically significant reduction in human MSC viability.^13^ Following fixation, the cells were imaged using confocal microscopy. The CPNs™ were readily taken up by the UC-MSCs, with the majority of cells within the population becoming labelled (Fig.1). The signal intensity of the cells labelled with the CPN™ 1000 nanoparticles appeared lower than that of the cells labelled with the other CPN™ probes and ICG. This was likely because the lasers and fluorescence filters available on the confocal microscope were not optimal for detecting these nanoprobes. The perinuclear staining pattern of the CPN™ probes is consistent with their accumulation in the endolysosomal compartment, which is typical for most cell labelling nanoprobes, including quantum dots^24^ and iron oxide nanoparticles.^25^ This was also the case for ICG, which is mainly taken up into cells via endocytosis.^26^ At the concentrations used, none of the CPN™ probes nor the ICG caused any noticeable effect on hUC-MSC viability.

### Flow cytometric analysis of hUC-MSCs labelled with CPN™ 770 nanoprobes or ICG

Prior to undertaking flow cytometry, we first assessed the radiant efficiency and photoacoustic signal intensity of the CPN™ nanoprobes *in vitro* using FI (IVIS Spectrum) and MSOT, respectively. Because the CPN™ 770 nanoprobe had the highest radiant efficiency and also the strongest signal with MSOT (Supplementary Fig.1), subsequent experiments were performed exclusively with this CPN™ nanoprobe. Flow cytometric analyses of the CPN™ 770 labelled hUC-MSCs was then undertaken to confirm that the majority of cells were labelled, and to determine the relationship between labelling concentration and fluorescence intensity. hUC-MSCs were labelled for 24h with the following dilutions of a stock concentration of 1×10^9^ CPN™ 770 nanoprobes/ml: 1:10, 1:20, 1:50, 1:100 and 1:200. Unlabelled hUC-MSCs served as a negative control, and ICG-labelled cells as the positive control. Even with the lowest concentration of CPN™ 770 nanoprobes, the majority of hUC-MSCs showed a noticeable increase in fluorescence compared to unlabelled controls, and the mean fluorescence intensity increased with increasing concentrations of nanoprobe; this was likely due to an increase in the number of nanoprobes per cell. The majority of ICG-labelled hUC-MSCs also showed a noticeable increase in fluorescence compared to unlabelled cells (Fig. 2). The concentration of ICG used to label the hUC-MSCs was 100 µg/ml over 30mins. Increasing the concentration of ICG would be unlikely to increase the signal intensity because at concentrations above 80 µg/ml, quenching starts to occur,^27^ and over 100 µg/ml, fluorescence intensity decreases sharply.^27^ This is because at higher concentrations, there is an increase in the ratio of ICG polymers compared to monomers, the former having a weaker yield of fluorescence.^27^ Moreover, higher concentrations would be expected to reduce the viability of the cells, as has previously been shown.^13^ In light of these earlier studies, the highest labelling concentration of ICG used here was 100 µg/ml.

### Fluorescence, MSOT and bioluminescence imaging of UC-MSCs labelled with CPN™ 770- or ICG following subcutaneous injection in mice

To compare the effectiveness of CPN™ 770 nanoprobes and ICG for tracking cells *in vivo* with FI and MSOT, hUC-MSCs were labelled with 1×10^8^ particles/ml (equivalent to 1:10 dilution of the stock) for 24h, or with 100 µg/ml ICG for 30 mins. hUC-MSCs expressing firefly luciferase (Fluc) were used for these experiments to establish if the labelled cells remained viable *in vivo*. Unlabelled Fluc^+^ hUC-MSCs and Fluc^+^ hUC-MSCs labelled with CPN™ 770 nanoprobes or ICG were injected into the dorsal flanks of mice at a concentration of 5×10^5^ cells in an injection volume of 100µl. Neat CPN™ 770 nanoprobes were also injected at a concentration of 1×10^8^ particles in 100µl to serve as a positive control (Fig. 3A). Mice were imaged shortly after administration using an IVIS Spectrum. Cells labelled with CPN™ 770 or ICG and the neat CPN™ 770 nanoprobes were readily visible and distinguishable after spectral unmixing (Fig. 3B). As expected, unlabelled control cells did not emit a detectable signal.

While under anaesthesia, the mice were imaged with MSOT and the signal was spectrally unmixed using the relevant spectra (Supplementary Fig. 2). Interestingly, although the signal intensity of the CPN™ 770- and ICG-labelled hUC-MSCs appeared similar with FI (Fig. 3B), with MSOT, the signal from the CPN™ 770-labelled cells and neat CPN™ 770 nanoprobes was noticeably stronger than from the ICG-labelled cells. In fact the ICG-labelled cells were not only barely visible, but also of the same intensity as other background signals seen at this wavelength (Fig. 3C and Supplementary Fig. 3). Prior to MSOT imaging, the mice were administered with luciferin to enable BLI to be performed immediately following MSOT while the mice were still anaesthetised. As expected, the unlabelled and labelled hUC-MSCs showed a detectable signal, indicating that the cells were viable, whereas no signal was detected from the neat CPN™ 770 nanoprobes (Fig. 3D).

When ICG is administered intravenously, it is mainly in the monomeric form and can be readily visualised in the vasculature using MSOT, a common application being the assessment of liver function.^28^ The reason why ICG-labelled hUC-MSCs gave only a weak signal in the current study is possibly because following endocytosis, the ICG becomes concentrated in the endolysosomal compartment, likely favouring an increase in the polymeric form of ICG that is known to have less favourable optical properties.^27^

Although the CPN™ 770 nanoprobes proved more effective than ICG for tracking cells with MSOT, the performance of the two contrast agents was comparable when used for tracking cells with FI. A possible explanation for this might be the larger number of photoactive units and larger absorption cross section in a conjugated polymer nanoparticle when compared to a single molecule imaging agent. This can result in a greater proportion of the absorbed light being converted to heat and ultrasound waves rather than being emitted as light, thereby generating a stronger signal in MSOT.^29^ Taken together, the results show that CPN™ 770 nanoprobes and ICG can both be used to track cells *in vivo* using FI, but the CPN™ 770 nanoprobes are far superior for tracking cells with MSOT.

## Conclusions

Here we assessed the potential of NIR CPN™ nanoprobes as cell tracking agents in comparison to the FDA-approved NIR dye, ICG. We found that similarly to ICG, nanoprobes with emission maxima ranging from 770nm to 1000nm were readily uptaken by hUC-MSCs, enabling the cells to be imaged *in vitro* using confocal microscopy. Following subcutaneous administration into mice, CPN 770™ -labelled hUC-MSCs and ICG-labelled hUC-MSCs could both be readily detected *in vivo* using FI. However, using MSOT, in contrast to CPN 770™ -labelled cells which were easily visible, ICG-labelled cells could barely be detected. NIR CPN™ nanoprobes, and CPN 770™ in particular, have great potential for cell tracking applications *in vivo* using FI and MSOT. The presence of iron oxide nanoparticles within the core of the nanoparticles also means that they could be useful for magnetic resonance imaging and magnetic particle imaging. Another advantage of CPN 770™ over ICG is that with the former, it is possible for the labelled cells to be analysed using microscopy following animal sacrifice. This would be difficult with ICG-labelled cells due to the rapid photodegradation of ICG.

## Author Contributions

Ana Muñiz-García: methodology, data collection and analysis, writing – original draft, writing – review and editing. Alejandra H Pichardo: methodology, data collection and analysis, writing – original draft, writing – review and editing. James Littlewood: methodology, data collection and analysis, writing – original draft, writing – review and editing. Jack Sharkey: methodology, data collection and analysis, writing – review and editing. Bettina Wilm: writing – review and editing, supervision. Hannah Peace: methodology, writing – review and editing. Dermott O’Callaghan: resources, methodology, writing – review and editing. Mark Green: conceptualization, data analysis, writing – original draft, writing – review and editing. Arthur Taylor: methodology, data collection and analysis, writing – original draft, writing – review and editing, supervision. Patricia Murray: conceptualization, data analysis, writing – original draft, writing – review and editing, supervision, funding acquisition.

## Acknowledgements

We gratefully acknowledge the support of the University of Liverpool’s Flow Cytometry Facility, Centre for Cell Imaging and Centre for Preclinical Imaging. This work was funded by a Business Interaction Voucher from the UK’s Royal Microscopy Society and the Biotechnology and Biological Sciences Research Council awarded to PM and Stream Bio, and by the European Union’s Horizon 2020 research and innovation programme under the Marie Sklodowska-Curie grant agreement No. 813839.

