## Supplementary material for "Near infrared conjugated polymer nanoparticles (CPN™) for tracking cells using fluorescence and optoacoustic imaging": Supplemetal Figs 1 to 3

### Slide 1
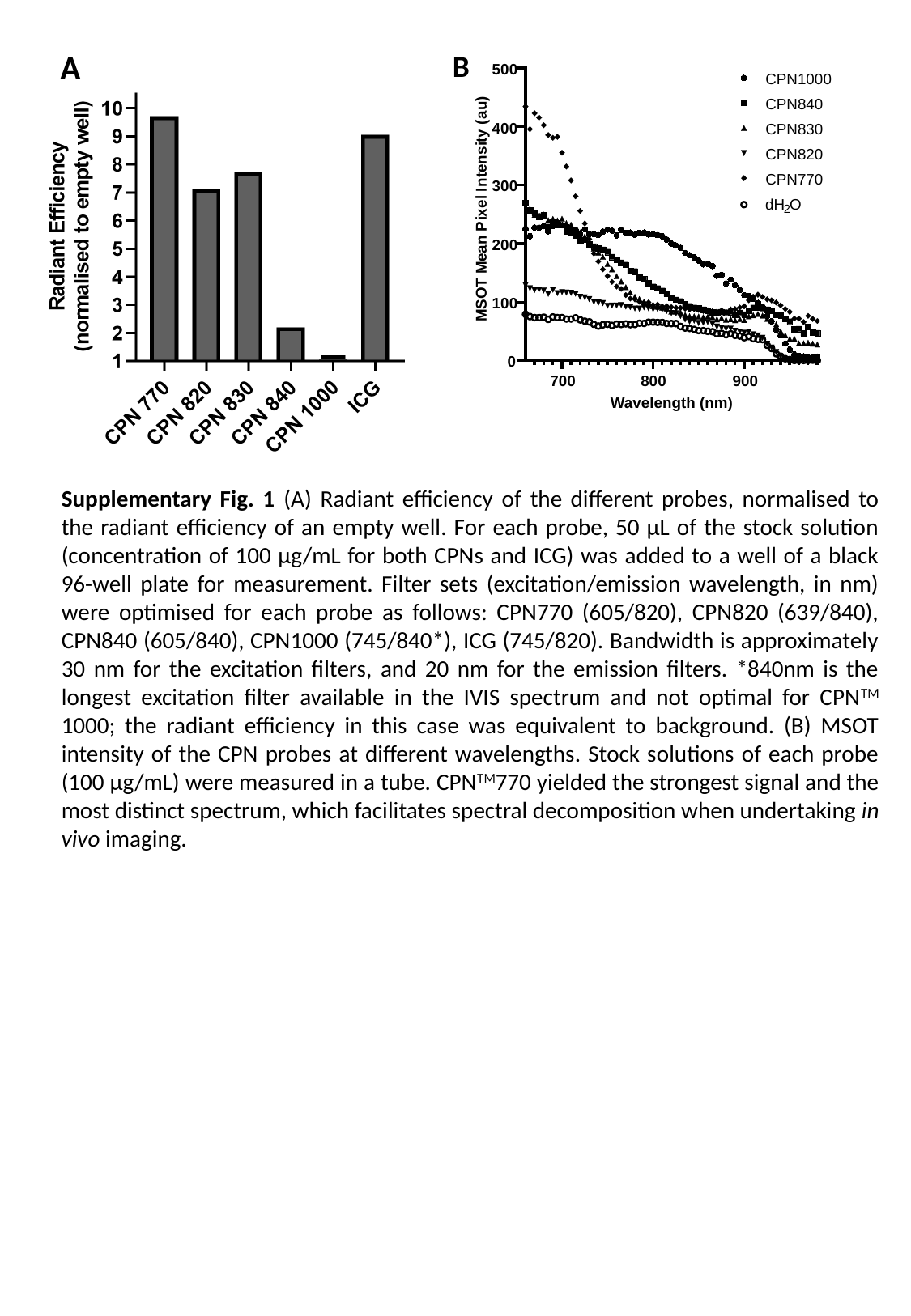

A
B
500
400
300
200
100
0
700
800
900
)
u
a
(
y
t
i
s
n
e
t
n
I
l
e
x
i
P
n
a
e
M
T
O
S
M
Wavelength (nm)
CPN1000
CPN840
CPN830
CPN820
CPN770
dH
O
2
Supplementary Fig. 1 (A) Radiant efficiency of the different probes, normalised to the radiant efficiency of an empty well. For each probe, 50 µL of the stock solution (concentration of 100 µg/mL for both CPNs and ICG) was added to a well of a black 96-well plate for measurement. Filter sets (excitation/emission wavelength, in nm) were optimised for each probe as follows: CPN770 (605/820), CPN820 (639/840), CPN840 (605/840), CPN1000 (745/840*), ICG (745/820). Bandwidth is approximately 30 nm for the excitation filters, and 20 nm for the emission filters. *840nm is the longest excitation filter available in the IVIS spectrum and not optimal for CPN^TM^ 1000; the radiant efficiency in this case was equivalent to background. (B) MSOT intensity of the CPN probes at different wavelengths. Stock solutions of each probe (100 µg/mL) were measured in a tube. CPN^TM^770 yielded the strongest signal and the most distinct spectrum, which facilitates spectral decomposition when undertaking in vivo imaging.

### Slide 2
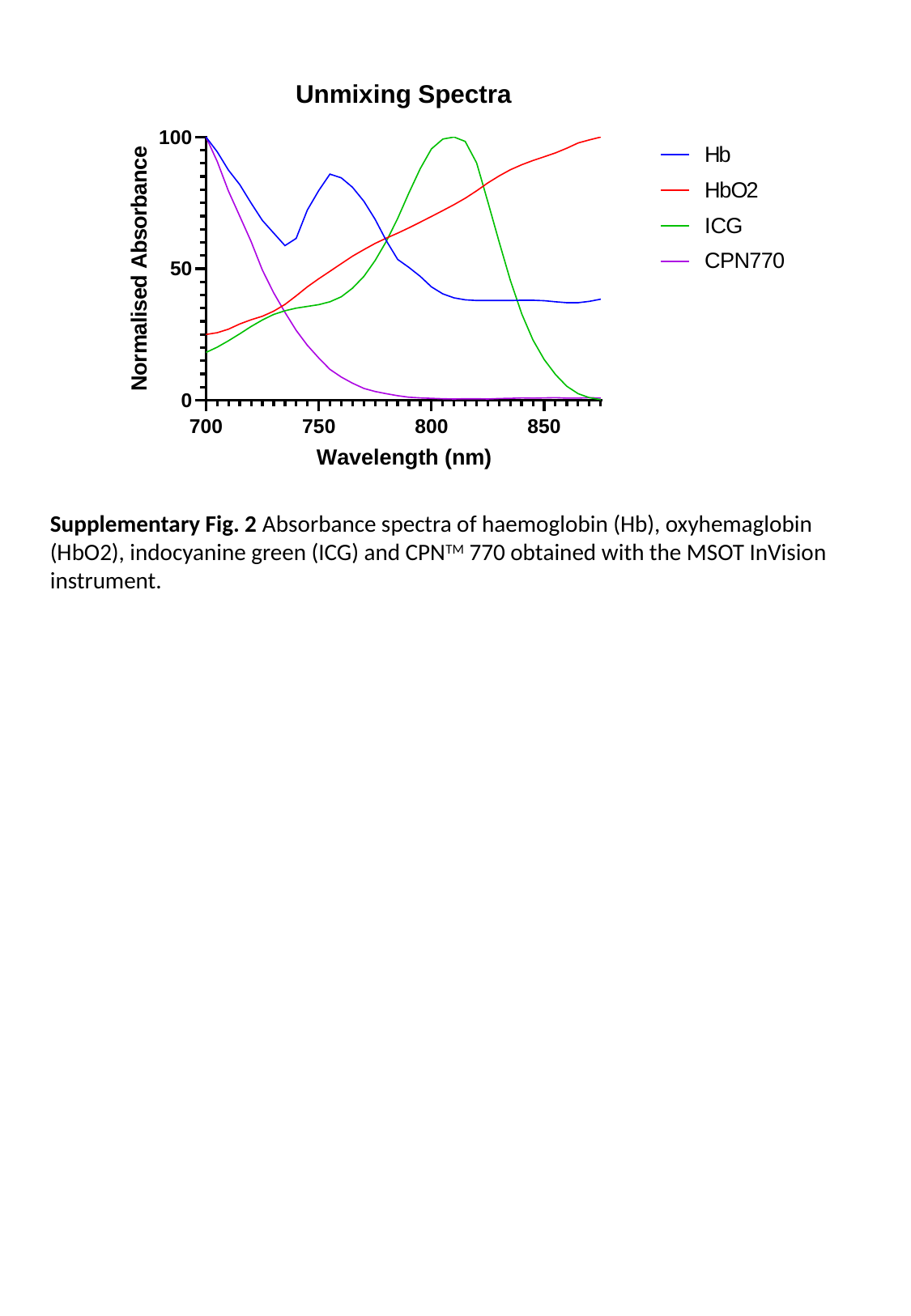

Supplementary Fig. 2 Absorbance spectra of haemoglobin (Hb), oxyhemaglobin (HbO2), indocyanine green (ICG) and CPN^TM^ 770 obtained with the MSOT InVision instrument.

### Slide 3
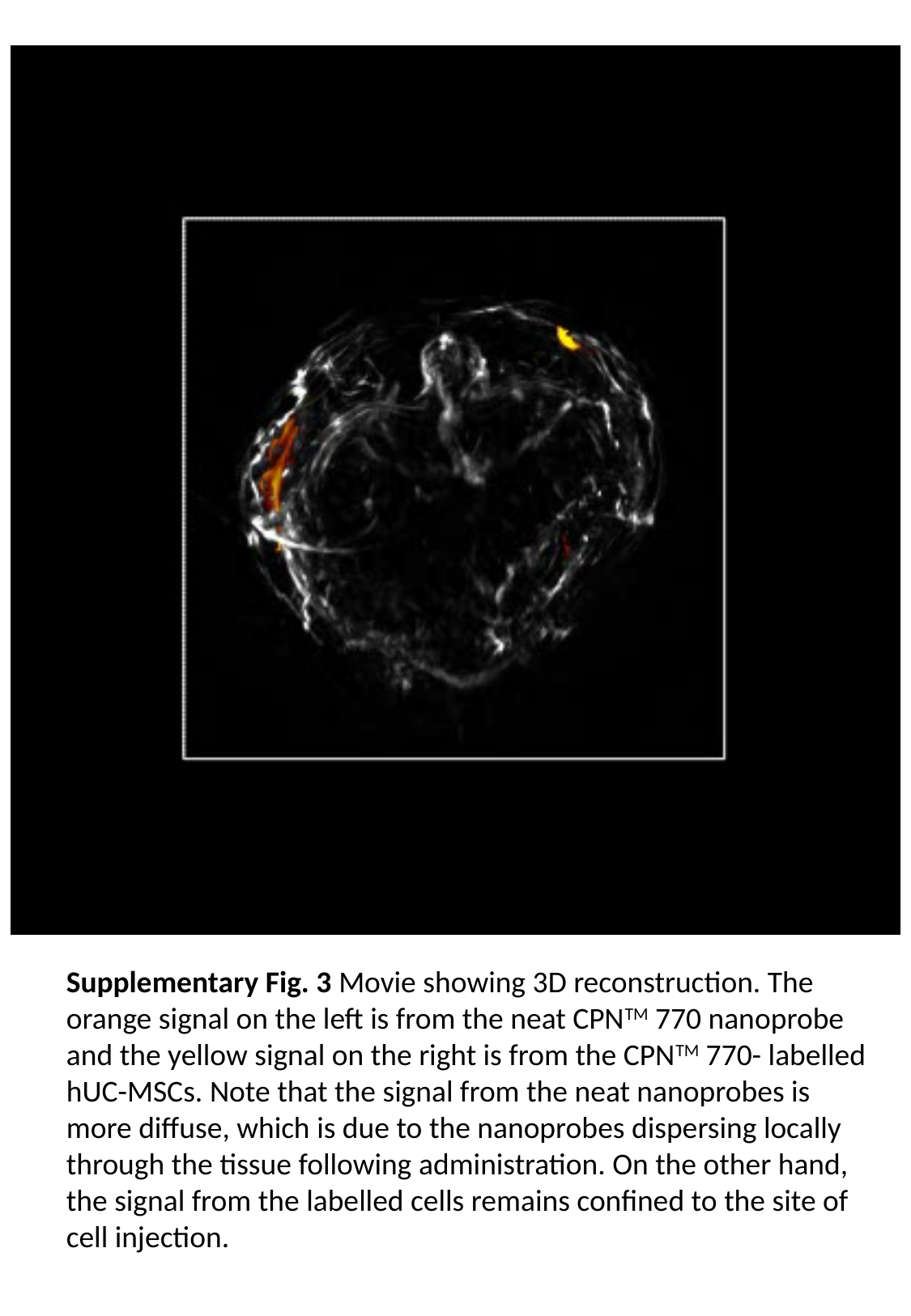

Supplementary Fig. 3 Movie showing 3D reconstruction. The orange signal on the left is from the neat CPN^TM^ 770 nanoprobe and the yellow signal on the right is from the CPN^TM^ 770- labelled hUC-MSCs. Note that the signal from the neat nanoprobes is more diffuse, which is due to the nanoprobes dispersing locally through the tissue following administration. On the other hand, the signal from the labelled cells remains confined to the site of cell injection.
